# Quantum Approximated Graph Cutting: A Rapid Replacement for T-REMD?

**DOI:** 10.1101/2020.12.11.420968

**Authors:** Samarth Sandeep, Sona Aramyan, Armen H. Poghosyan, Vaibhav Gupta

## Abstract

Determining an optimal protein configuration for the employment of protein binding analysis as completed by Temperature based Replica Exchange Molecular Dynamics (T-REMD) is an important process in the accurate depiction of a protein’s behavior in different solvent environments, especially when determining a protein’s top binding sites for use in protein-ligand and protein-protein docking studies. However, the completion of this analysis, which pushes out top binding sites through configurational changes, is an polynomial-state computational problem that can take multiple hours to compute, even on the fastest supercomputers. In this study, we aim to determine if graph cutting provide approximated solutions the MaxCut problem can be used as a method to provide similar results to T-REMD in the determination of top binding sites of Surfactant Protein A (SP-A) for binding analysis. Additionally, we utilize a quantum-hybrid algorithm within Iff Technology’s Polar+ package using an actual quantum processor unit (QPU), an implementation of Polar+ using an emulated QPU, or Quantum Abstract Machine (QAM), on a large scale classical computing device, and an implementation of a classical MaxCut algorithm on a supercomputer in order to determine the types of advantages that can be gained through using a quantum computing device for this problem, or even using quantum algorithms on a classical device. This study found that Polar+ provides a dramatic speedup over a classical implementation of a MaxCut approximation algorithm or the use of GROMACS T-REMD, and produces viable results, in both its QPU and QAM implementations. However, the lack of direct configurational changes carried out onto the structure of SP-A after the use of graph cutting methods produces different final binding results than those produced by GROMACS T-REMD. Thus, further work needs to be completed into translating quantum-based probabilities into configurational changes based on a variety of noise conditions to better determine the accuracy advantage that quantum algorithms and quantum devices can provide in the near future.

## Introduction

Determining the most optimal protein configurations for binding models is one of the most important tasks to calculate before completing protein-protein, protein-ligand, or further inter-protein analysis. The complex surface characteristics of proteins, even in their crystallographic form without solvent interactions as found within Protein Data Bank (PDB) formatted files, means that the complete computation of binding characteristics of entire structures can be anywhere from infeasible [29] to impossible on classical computers [7] without model simplification or cutting the number of residues involved in modeling based on experimental results. While determining the most optimal residues for binding models is a straightforward process if experimental data exists for binding kinetics of that protein, it is difficult to complete intuitively for proteins of which there is little known experimental binding behavior, as chemical formation kinetics is completed in non-deterministic, polynomial (NP) time on a classical device [7].

To solve this problem, a multitude of bioinformaticians have turned to tools that can complete molecular dynamics simulations of the proteins undergoing thermal relaxation in order to more effectively reveal their best binding sites. While tools such as Rosetta [14] provide this functionality in the form of *ab initio* modeling of protein energies and create rigid protein models, Schrodinger [21] and GROMACS’ Temperature-Replica Exchange Molecular Dynamics (T-REMD) [38] complete this task by completing iterative molecular dynamics simulations of each of the protein characteristics’ interactions.

However, the completion of these approximated computational models can be quite time consuming and computationally complex themselves [10]. With a maximum GROMACS performance of 304 ns/day achieved across 600 physical cores with 2 GPUs used per core of access [5], the scale of computational resources needed for molecular dynamics simulations for even multi-hour levels of full protein analysis means that GROMACS is still quite impractical to use at scale when analyzing full interactomes or virions [27].

Consequently, there remains a need to complete the protein binding site analysis before docking at the precision level of molecular dynamics with much faster speeds and with lower cost computational systems to dramatically increase the ability to computationally model full interactomes in order to effectively model protein-protein and protein-ligand binding at scale.

### Topological Analysis of Protein Surfaces for Binding Site Analysis

Recently, there has been a growing amount of work focused on finding the minimal and maximal points of protein models through the analysis of protein shapes, or topology, before adding electrostatically- or electrodynamically-based equations for full pre-binding analysis in order to screen for top sites prior to compute-intensive force calculation. The completion of this task on classical computers with clever, third-degree polynomial topological algorithms [6] or by quantum annealing devices that straddle the line between highly optimized classical computing devices and fully quantum computational devices within a first-degree polynomial complexity class [25] has led to a new paradigm within biological modeling that is focused more on topological feature distinction.

Due to the realization of the benefits that could be gained by completing topological models to the overall time of computation, it seemed that the abstraction could be taken one step further through the implementation of Maximum Cut, or MaxCut, graphing algorithms [17]. Not only would these algorithms be able to effectively find the minima and maxima on protein surfaces, but they could actively apply a cost function across the interaction points between the different atoms within the protein to more effectively pick the best binding site with regards to electrodynamics and electrostatics as well. But, while there are potential estimations of a potential Maximum Cut that exist in various algorithmic forms, such as Greedy Cut [39] and the Goemans-Williamson algorithm [19], it has not been formally solved at scale, making it potentially infeasible for protein structure analysis.

### Quantum Computing in Improving MaxCut’s Odds

Quantum processors are devices that use the effects of quantum mechanics for methods of information transfer among bit-like devices. Due to this ability for the bits themselves to hold multiple states, quantum computers seem like ideal solver devices for molecular dynamics calculations, as they could represent every electron within a protein. However, current quantum processors are still in their infancy; the largest superconducting quantum processor that exists as per the writing of this paper is the Google Bristlecone device, with 72 qubits [8]. As 1 qubit is essentially 1 extra large electron [22], it would take hundreds of qubits to model even the smallest of proteins.

However, due to their ability to provide superposition and entanglement features on bits, quantum computers have the potential to take exponential scale problems and turn them into polynomial or even log scale problems [4]. As such, they could be devices that can potentially provide massive improvements to a MaxCut problem. Particularly, the quantum approximate optimization algorithm (QAOA) has been found to be a potential solution to this problem [15]. This hybrid quantum-classical algorithm combines the quantum computer’s ability to effectively solve exponential problems with a classical cost function to determine the best cuts within a set of quantum bits, and as such, could provide benefits with few qubits if cleverly designed to scale effectively with classical devices. As such, QAOA has been seen to have biological uses in the past, with the team at ProteinQure using QAOA to effectively fold small proteins [16]. Thus, QAOA seems to be a good candidate for potentially replacing T-REMD.

### Polar+: Suitable competition for Goemans and Williamson and GROMACS?

For the past eight months, Iff Technologies has been working with academic and startup clients to test its software, Polar+ [33]. Polar+’s protein pruning tool is an implementation of the QAOA algorithm that includes further bioinformatics contextualization. It has been shown to improve the binding outcomes of protein-ligand interactions [34], However, it has not been directly compared to GROMACS T-REMD and Goemans-Williamson implementations at scale until this point.

In this paper, Polar+’s protein pruning tool running on a quantum computer connected to a 1-node classical computer is compared in performance to an instance of Polar+’s protein pruning tool running on a quantum abstract machine running on the Pittsburgh Supercomputing Center’s Bridges’ Supercomputer, a Goemans-Williamson implementation running on Bridges’ Supercomputer at the Pittsburgh Supercomputing Center, and to a GROMACS T-REMD implementation running on the Bridges’ Supercomputer at Pittsburgh Supercomputing Center and the JUWELS Module 1 Supercomputer at the Forschungszentrum Jülich, North Rhine – Westaphalia, Germany, for the binding site analysis of Surfactant Protein A (SP-A), an epithelial membrane collectin that could have immune protective properties against COVID-19. This study aims to answer two major questions:

1. Can graph cutting using MaxCut estimators effectively emulate GROMACS T-REMD molecular dynamics studies used before docking studies?
2. Does Polar+ running on a quantum processing unit provide true quantum advantage over a classical implementation of the quantum algorithm and Goemans-Williamson?

## Materials and Methods

### Biomolecular Structures

The crystallographic coordinates for the SP-A protein structure were determined using Uniprot [13]. Through analysis of this database, the model 5FFR [20] was determined to be the most all-encompassing structural model of SP-A, covering 147 of its amino acids at a resolution of 2.20 Å. However, this model did include phosphocholine ligands, which could interfere with direct interrogation of the amino acids that make up the structure of SP-A [20]. Thus, the phosphocholine was removed from the 5FFR model before binding site analysis.

### T-REMD with GROMACS

For the completion of the T-REMD analysis, GROMACS 5.0.4 was utilized to create a suitable solvent environment, along with a set of temperature and pressure controls, in order to most accurately determine the protein configuration in a binding environment, as completed in similar studies [35]. Being a molecular dynamics software, GROMACS completes sets of multi-axial nearest neighbor calculations for a set of forces for coordinate position and velocities across a number of time steps [9]. First, forces for each molecule within a solute and solvent are calculated using a prescribed set of forces unique to different solvent environments. As this study aims to understand a protein in a neutral solvent environment, three different force models were used: the Assisted Model Building with Energy Refinement (AMBER) force field, the Optimized Potentials for Liquid Simulations (OPLS) force field, and the CHemistry At Harvard Macromolecular Mechanics(CHARMM) force field.

Being the oldest force field used, AMBER [11] has the simplest form, with total potential energy for a macromolecule following a summation between bond energy as an ideal spring, geometrical energy from each angle within the covalent bonding between atoms, torsioning due to bond order, and intra-atomic forces represented as a van der Waals force added to an electrostatic force, wherein *f_ij_* represents the Fourier transformation, *E_ij_* represents the well depth of the atom’s location, and other constants represent their respective parts. This study used AMBER99sb.

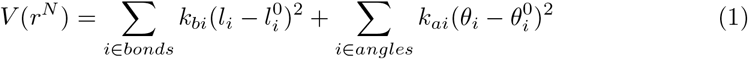

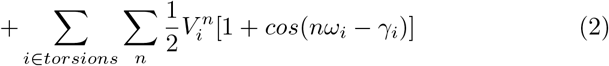

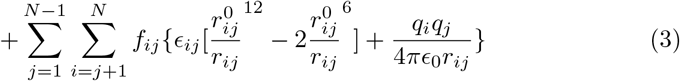

**Equation 1**. AMBER Force Field Formula adapted from Case *et al*. for Electrical Potential Across Protein [11]

OPLS [23] shares much of the same structure as AMBER. However, it aims to provide better analysis of the differences between bonded, nonbonded, and dihedral atoms, which are present on a multitude of energetic planes, through the use of torsional and electrostatic constants derived for each element and each organic functional group, represented as A and C. OPLS is also designed for use with the TI3P water model, which is a 3-side rigid water molecule with charges, as the default solvent for the force.

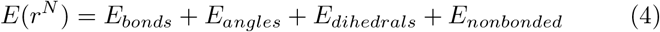

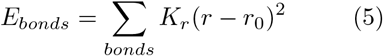

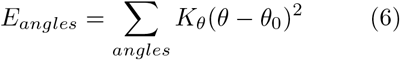

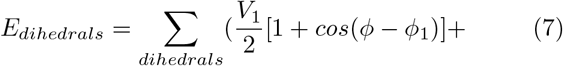

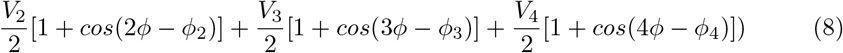

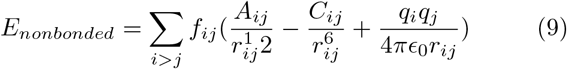

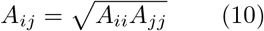

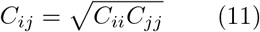

**Equation 2.** OPLS Force Field Formula adapted from Jorgensen et. al. for Electrical Field Across Protein [23]

CHARMM [24] is a force-field model that aims to take OPLS further through the addition of an impropers and a Urey – Bradley term, which intend to improve upon the torsional modeling of the atomic interactions through the accounting of bending and non-binding interactions between atoms in the 1,3 positions of an organic molecule due to proximity of electrostatic forces, respectively. This study used CHARMM36.

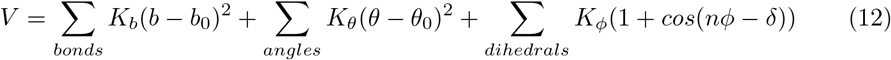

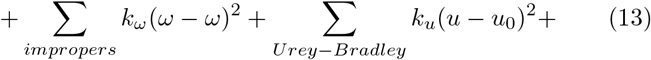

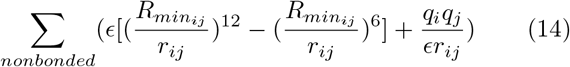

**Equation 3.** CHARMM force field formula adapted from Mackerell *et al*. for electrical potential across protein [23]

The following system has been utilized for the benchmarking of the GROMACS software:

SP-A protein in water bulk consisting of ~62,000 atoms (number of water molecules ~20,000) in 6.5×6.5×6.5 nm^3^ cell.

The testing runs were done using the following parameters: 2fs timestep, PME electrostatics, and van der Waals forces truncated at 1.2 nm with corresponding pressure and temperature control. The benchmark runs were typically for 10000 steps (20ps) with/without writing output any trajectory and coordinate files (Note that with no write trajectories and confout slightly increases the performance). For our tests, we use the “-pin on” and “-dlb yes” GROMACS flags, where “-pin on” stops the kernel from moving processes between cores by locking the cores, and allow dynamic load balancing to automatically run when the load imbalance is 5% or more, which is important for inhomogeneous systems. Note that for optimal performance, we also try mdrun −resethway and −maxh=0.05 options, which corrects the benching results. After these first test runs, the force fields for SP-A were taken into consideration for a total of 10 ns, or 5,000,000 time steps, in order to be able to obtain reasonable accuracy with regards to the interaction of SP-A within a water model.

### T-REMD Device: JUWELS Supercomputer

JUWELS multi-petaflop supercomputer [36] located at the Julich Supercomputing Centre (JSC Germany) is one of the most powerful computing resources available in Europe. It consists of a total 2567 compute nodes (2511 CPU-only partitions and 56 Nvidia V100 GPU nodes), where the nodes are interconnected through Mellanox Infiniband high performance network architecture. The CPU-nodes are equipped with two Intel Xeon Platinum 8168 processors (base frequency of 2.7GHz), while GPU-nodes are fitted with the two 2.4GHz Intel Xeon Gold 6148 processors. Each GPU node contains four Nvidia V100 cards with 5120 CUDA cores. Note that the peak performance of the mentioned cluster is ~4,15 TF/s based on the Linpack Benchmark.

### T-REMD and Classical Graph Cutting Device: Bridges at the Pittsburgh Supercomputing Center

The Bridges Supercomputer at the Pittsburgh Supercomputer Center has 752 Regular Shared Memory (RSM) nodes, each consisting of 2 Intel Haskell CPUs with 14 cores per CPU, as well as 9 AI-GPU nodes, each consisting of 2 Intel Xeon Gold 6148 CPUs with 20 cores each and 8 NVIDIA Volta V100 GPUs. Due to GROMACS’ ability to provide improvements to performance through the use of GPUs, the AI-GPU nodes were used for the completion of OPLS, CHARMM, and AMBER T-REMD analyses on SP-A. Additionally, these nodes were utilized for the completion of the Goemans-Williamson interpretation of the MaxCut problem.

### Goemans-Williamson Implementation

The Goemans-Williamson algorithm was implemented through the use of the CVX Graph Algorithms Python package [1] across the entire atom map of the protein. In this implementation of the algorithm, the atoms that were identified to be cut by the algorithm were cut from the map, leaving a map of what should be the most energy-resilient atoms, and therefore the key binding sites on the protein.

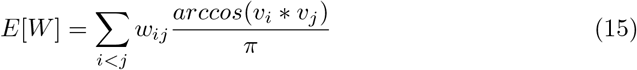

**Equation 4.** Goemans-Williamson MaxCut algorithm [19], where *E[W]* represents the expectation value of a node, *i* and *j* representing the two dimensions of node movement, *w* representing the weight of each node, and *v* representing the vector the node produces itself

In order to run this algorithm as effectively as possible, it was made to run on the Bridges Supercomputing System with PySpark [2] used as the batching mechanism between nodes. Besides this addition, there were no more changes made to the CVXGraph Goemans-Williamson algorithm used.

### Polar+ Protein Pruning Implementation

Using the quantum computer as analogs for the atoms in the proteins, sets of 3 atoms each were placed on qubits next to each other in placements that were topologically similar to the interaction space between the atoms themselves as identified by their PDB files. Then, the QAOA MaxCut package from Rigetti and Co. [3] was utilized to implement the MaxCut process on the 3 atom subgraph of those qubit positions on the Rigetti Aspen 8 quantum computer.

The Rigetti Aspen 8 is a quantum computing device that operates using superconducting Josephson Junctions to create a silicon based lattice structure of 31 qubits, or quantum bits, are embedded onto a piece of gold and cooled to nearly 0°K through the use of helium based cooling chambers [31].

Similar to the Goemans-Williamson implementation, the QAOA approach to MaxCut is a summation across a set of different weights posed by the vectors in the graph to gain an expectation value for the cut size. However, rather than using weights either provided by or produced arbitrarily based upon the presumed magnitude of each vector, the weights used here are the tensor product of the lowest eigenvectors of each qubit’s x-axis operators, *σ_x_*.

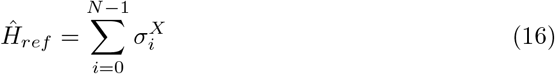

**Equation 5.** Total Energy of Each Operator within a Quantum Circuit of N Nodes as per the Rigetti Implementation of QAOA [3]

Using the qubits and their angular rotations themselves, which is determined after running the algorithm over a set of test runs for a particular use case and determining which results provide the most problem-specific information, as the nodes within the graphs that MaxCut must be performed on, the total energy of each of these operators simply needs to be decomposed to determine to which node the maximum cut value belongs to. This can be completed using the Trotter-Suzuki algorithm [3] for each angle of rotation. Due to the ability for the *RX* and *RZ* gates of the Rigetti quantum processor set to create these deconstructed values, these gates simply need to be run for the angles used for each node.

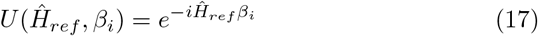

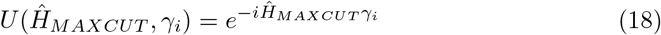

**Equation 6.** Potential Energy Found at Each Qubit after Decomposition According to the Trotter-Suzuki Algorithm [3]

At the end of this process, basis states representing different qubits being cut from the graph were provided at different probability levels. These basis states, which were binary numbers in order to the qubit and flip state of that qubit, were contextualized to refer to the qubit that needed to be cut from the graph; 1s were taken to be qubits that were part of the new graph, and 0s were taken to be qubits that were eliminated. To find the best binding site in the best configuration, then, the highest probability basis state was evaluated, and the atoms with positions that had 0 values within the basis states calculated were eliminated from the overall list of protein atom positions.

Lastly, these atom positions were then cross referenced to the atoms they originally referred to in order to determine which atoms need to be part of a new PDB file representing only the best binding sites. Finally, this conversion took place using the Biopython software package.

### Measuring effectiveness of models through docking using ZDOCK

In order to determine the effectiveness of each model in determining top binding sites, the models were each made to bind with the SARS-CoV-2 spike protein in its open conformation (PDB: 6VSB). The binding was completed through the use of the software ZDOCK, created by the University of Massachusetts Medical Center [12]. Within this software, top binding sites are determined through the closeness of a summation of Fourier functions of topological and desolvation energetic parameter scalars, as well as electrostatic values from CHARMM, for each atom in 6 dimensions.

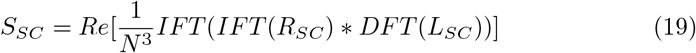

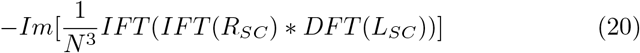

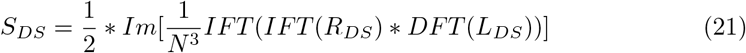

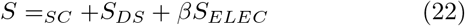

**Equation 7.** Calculation of scalar energetic from the aggregation of shape complementarity(SC) for the larger protein, the receptor (Rsc), and the smaller protein, the ligand (Lsc), as well as the desolvation energy for derived through the summation of each contact potential atomic contact point’s desolvation energy obtained experimentally from Zhang *et al*. [40], wherein *α* and *β* are scaling ratios used to ensure that *S_sc_* and *S_elec_* are of the same scale.

Because of the binding capability being reflected as a scalar, a higher *S_total_* value reflects stronger binding value. Using the top 2000 conformations produced by ZDOCK for the AMBER, OPLS, CHARMM, Goemans-Williamson, and Polar+ models used in this study, the top binding candidates can be reviewed against other SP-A results, and the Root Mean Square Deviation (RMSD) across the top 2000 conformations can be calculated to determine which model produces the highest accuracy conformations through the use of ZDOCK.

## Results

### T-REMD on JUWELS: Computational Performance

In the table below, GROMACS performance on JUWELS supermachine as a function of cores/logical cores is shown. The mentioned benchmark was done in as follows: we added #SBATCH − -nodes=N (N=1-6) and #SBATCH − -ntasks-per-node=96 lines to a SLURM script, which helps to distribute the job across all nodes available. As already mentioned, the performance was carried out without manipulating the OpenMP thread per MPI process parameter. The curve plotted in the following figure show that increasing the number of cores/logical cores leads to the increase of performance until a certain point. Thus, we track that the parallelization scales well up to 192 cores, as further cores strain communication within the GROMACS program. To achieve better performance of GROMACS for a single node and check the OpenMP efficiency, we have done some tests, where tasks were controlled by SLURM –ntasks option and environmental variable – OMP_NUM_THREADS, and the threads were controlled using mdrun −ntomp flag. We identified that altering OpenMP threads slightly changed the performance; however, in overall, the “threading effect” does not give any significant changes when using a single node.

**Table 1.** Performance Benchmarking for 10000 steps on a Single Node on JUWELS

| MPI Processes x OpenMP Threads | Performance (ns/day) |
| --- | --- |
| 16 x 6 | 72.558 |
| 24 x 4 | 80.368 |
| 32 x 3 | 79.776 |
| 48 x 2 | 79.840 |
| 96 x 1 | 85.327 |

Multi-node “threading” experiments showed that the performance of 62K system depends on threads/processes ratio. We performed a series of tests by changing the threads/processes ratio when four nodes are selected. The performance and average load imbalance results are shown in Table 2. We saw that further increase of number of threads led to the decrease of performance and correspondingly, the increase of unbalanced part of the molecular dynamics study.

**Table 2.** Performance Benchmarking for 10000 steps with Average Load Imbalance on a Four Node Setup on JUWELS

| Processes / Threads | Performance (ns/day) / Average Load Imbalance (%) |
| --- | --- |
| 384/1 / 6 | 264.674 / 8.7 |
| 192 / 2 | 242.593 / 9.5 |
| 128 / 3 | 204.361 / 13.9 |
| 96 / 4 | 192.470 / 17.7 |
| 64 / 6 | 183.105 / 19.1 |

**Table 3.** Peformance data for each force field using 10,000 step benchmark on both devices

| Device Used | Force Field | Number of Atoms | Number of Waters | Average performance (ns/day) | Average Wall Time (s) |
| --- | --- | --- | --- | --- | --- |
| JUWELS 48 Cores (96 Logical) | CHARMM | 61896 | 19863 | 47.652 | 18.135 |
| Bridges 28 Cores with Four NVIDIA Tesla K80s | CHARMM | 61896 | 19863 | 104.05 | 8.3056 |
| JUWELS 48 Cores (96 Logical) | OPLS | 60197 | 19310 | 108.25 | 7.9830 |
| Bridges 28 Cores with Four NVIDIA Tesla K80s | OPLS | 60197 | 19310 | 111.619 | 7.7420 |
| JUWELS 48 Cores (96 Logical) | AMBER | 60191 | 19308 | 108.38 | 7.9730 |
| Bridges 28 Cores with Four NVIDIA Tesla K80s | AMBER | 60191 | 19308 | 108.235 | 7.9840 |

**Table 4.**
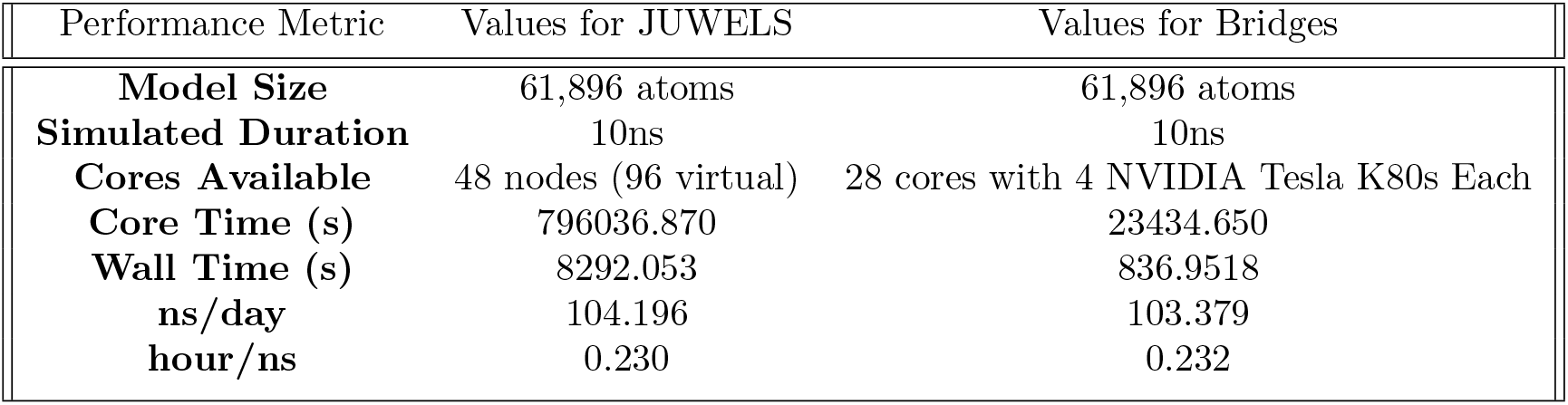
Completion Time for CHARMM Model on Both Devices for 5000000 steps

**Table 5.**
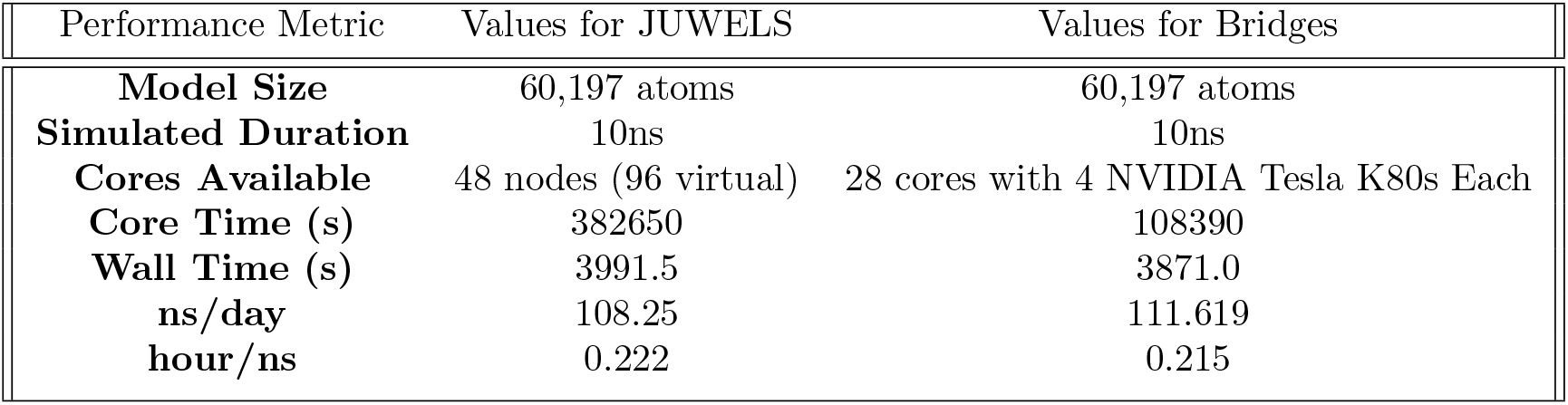
Completion Time for OPLS Model on Both Devices for 5000000 steps

**Table 6.** Completion Time for AMBER Model on Both Devices for 5000000 steps

| Performance Metric | Values for JUWELS | Values for Bridges |
| --- | --- | --- |
| **Model Size** | 60,191 atoms | 60,191 atoms |
| **Simulated Duration** | 10ns | 10ns |
| **Cores Available** | 48 nodes (96 virtual) | 28 cores with 4 NVIDIA Tesla K80s Each |
| **Core Time (s)** | 383116 | 111779 |
| **Wall Time (s)** | 3986.5 | 3992.0 |
| **ns/day** | 108.38 | 108.235 |
| **hour/ns** | 0.221 | 0.222 |

**Table 7.** Average Completion Time for Goemans -illiamson Model And Vector Outputs over Four Trials

| Performance Metric | Value |
| --- | --- |
| **Time to Completion (s)** | 11417.92 |
| **Number of Input Vectors** | 2284 |
| **Number of Output Vectors** | 540 |

**Table 8.** Time Completed and Vectors Processed for Each Implementation of the Polar+ Protein Pruning Technology

| System Used | Number of Qubits Used in Each iteration | Number of Qubits in Quantum System | Time Taken To Completion (s) |
| --- | --- | --- | --- |
| Rigetti Aspen 8 QPU | 3 | 31 | 528.68 |
| Rigetti QAM Operating on 32 CPU Instance on Digital Ocean | 3 | 29 | 384.84 |

Next, this same benchmark of 10,000 steps was completed for the completion of each force field on both the Bridges and JUWELS devices under constant Number, constant Pressure, constant Temperature (NPT), or isobaric, conditions, as shown in the tables below.

With regards to the actual implementation for each of the force fields, only 48 cores were available for running 5,000,000 time steps. As such, the performance obtained was less than that observed in the test runs. This led to a fastest observed run time of around 1 hour and 6 minutes for the AMBER model on JUWELS, and around a 14 minute run time of CHARMM on Bridges. Below is the data for the each model on both devices.

### Goemans-Williamsons Implementation: Computational Performance

The time taken for total completion of the Goemans-Williamson process for graph cutting across the whole protein, as well as the number of input and output vectors for each atom prior to and after the algorithmic implementation, are shown here.

### Implementation of Polar+ on both QPU and QAM: Computational Performance

The time taken for total completion of the Polar+ based protein pruning process for graph cutting across the whole protein are shown here for both the QAM and the QPU. Additionally, the number of qubits used and the total number of qubits of the system during algorithmic implementation are provided.

### Produced SP-A models from each software

The newly configured models from GROMACS and cut models from Polar+ and Goeamans-Williamson implementations can be found below. While the models produced by graph cutting methodologies do not have the same dramatic configurational changes as observed by the GROMACS models, all models have the same protruding regions, indicating that the most effective binding sites across all structures are similar. Looking at the GROMACS models further, it appears that the model produced by AMBER has relatively little change from the original SP-A model, while the models produced by OPLS and CHARMM are much more inwardly folded.

### Compared ZDOCK Docking to SARS-CoV-2 Spike Across All Models

The top docking score, docking pose, and Root Mean Square Deviation (RMSD) across docking poses for each of the five SP-A models can be found in the figure below. Based on these figures, we found that the docking poses created with graph cut models are statistically more significant than those found by GROMACS, and the sites of binding observed on SP-A are from the same residue across all models. However, while each of the GROMACS based models bind to the open receptor binding domain (RBD) of SARS-CoV-2 spike, each of the graph cutting based methods bind to the subunit 2, or the base, of the spike instead.

## Discussion

### Differences in Computational Complexity and Algorithmic Accuracy

As observed in the results, Polar+ was able to complete the task of top binding site determination in a time faster than that observed by GROMACS, and dramatically faster than that observed by the Goemans-Williamson algorithm, with the same range of Binding Sites found by OPLS, Polar+, and AMBER. However, the added atom-by-atom configurational change observed by GROMACS led to direct binding at the RBD of SARS-CoV-2 spike.

The reason for this increase in speed is the ability of the quantum processor to utilize superpositions of each state and entanglement between states to produce strong cut probabilities nearly instantaneously for multiple atoms, while classical processing devices must complete these tasks serially for each atom. This allows for a slimmer algorithmic approximation that does not require multi-axial force calculations or higher order functions. More importantly, the steps completed by GROMACS to properly characterize the forces for each atom are additive amongst the atoms and then additive amongst each pair of atoms, creating a computational complexity of *O*(*n*^2^) for each atom [10]. But, this still does not compare to the steps that need to be completed for Goemans-Williamson, which, while having a lower dimensionality requirement, has a complexity of *O*(*n*^2^*logn*) [28] due to its arccos term. Thus, QAOA is naturally the fastest algorithmic implementation, with an expected performance in the *O*(*logn*) regime [15].

Surprisingly, however, the QAM-based Polar+ implementation running on a medium-scale classical device was able to outperform the QPU based implementation. While this means that, in this interpretation, the QPU does not outperform a classical computing device for this task, it does mean that the procedural generation of quantum states on a classical device can provide algorithmic speedups to algorithms within the Bounded Quantum Polynomical, or BQP, regime, in which problems are of greater complexity than those that can be solved within deterministic Polynomial (P) time but lesser complexity than those that require Non-Polynomial deterministic time (NP) [26], thus providing further evidence of a quantum algorithm use case that can, if coupled with an exponential time problem on a quantum device, can provide a definite quantum advantage as raised by Ge and Dunjko [18].

With regards to accuracy of the configurational spaces, GROMACS is the only software used here that effectively managed to capture the effects of a saline solution on the protein in the form of direct coordinate shifts, along with the initialization of a water model to ensure neutral solvency. This process, which leans heavily on atom-by-atom calculation of Coulombic forces, was expected to scale exponentially as atoms were added due to each additional, squared Coulombic force if the same time steps were to be used for all atoms. However, the number of time steps was capped at 5,000,000 steps, which allowed for a P-time implementation.

### Binding Accuracy

In Table 9, one can see the comparison of main results across all the discussed models. Based on this table, the following points can be ascertained:

**Table 9.** Docking Statistics for Each Model, with the Polar+ Model Representing both the Goemans-Williamson and Polar+ Models

| Complexes | Spike binding site | SP-A binding site | Spike binding chain and resid | SP-A binding resid | ZDOCK affinity value | RMSD Avg Å |
| --- | --- | --- | --- | --- | --- | --- |
| **5FFR & 6VSB** | S1, partially include open RBD | C-type lectin domain & Collagen like domain | C; GLN-493<br>A; ASN-440<br>C; ASN-487 | TYR-92<br>ASN-141<br>SME-111 | 1736.984 | 141.4112 |
| **AMBER SP-A & 6VSB** | S1, open RBD | C-type lectin domain & Collagen like domain | A; TYR-421<br>A; ALA-352 | SER-187<br>TRP-191 | 2017.295 | 150.6609 |
| **CHARMM SP-A & 6VSB** | S1, partially include open RBD | Collagen like domain | C; TYR-453<br>A; PRO-384 | GLN-97<br>TYR-92 | 1656.588 | 173.1233 |
| **OPLS SP-A & 6VSB** | S1 | C-type lectin domain | C; ALA-372<br>B; TYR-489 | GLN-108<br>SER-156 | 1817.81 | 133.9383 |
| **Polar+ SP-A & 6VSB** | S1 | C-type lectin domain | B; ASN-370<br>B; NAG-1317 | ARG-146<br>LYS-159 | 1333.335 | 219.7738 |
| **Polar+ SP-A & Polar+ Spike** | S2 | C-type lectin domain | C; NAG-1301 | TYR-188 | 448.202 | 43.5559 |

#### 1. All Binding locations as found by ZDOCK represent bindable regions

It can be seen that all of the models produced by GROMACS T-REMD, regardless of force field used, represents a binding site on SARS-CoV-2 Spike’s S1 segment in some cases partially including open Receptor Binding Domain (RBD) (chain A). Only the models consisting of proteins processed with Polar+ and Goemans-Williamson show binding onto the S2 segment of Spike. Additionally, it is worth mentioning that this model shows purely N-acetyl-*β*-glucosaminide (NAG) residues being involved in binding from Spike’s side. This model shows that the C-type lectin domain, with a N-linked glycosylation site, is involved in binding from SP-A’s side, which can allow for glycosylated bond formation. From literature, it is seen that the C-type lectin domain is responsible for carbohydrate recognition (see Figure 1) and NAG residues provide the glycosylation of Spike protein that are mainly present on the S2 segment, leaving the RBD open for binding to receptor [30]. Therefore, while the binding model produced by graph cutting methods make more sense when looking at the exact binding interaction from the perspective of SP-A, where there is a NAG residue interacting with an N-linked glycosylation site, the binding models produced by T-REMD make more sense from the perspective of SARS-CoV-2, as strongest binding is more likely to take place within the RBD.

**Figure 1.**
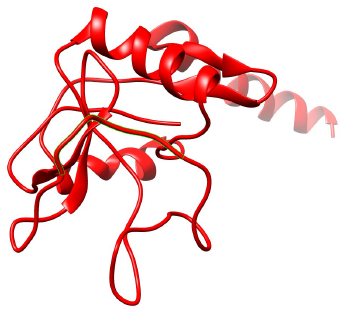
Model of SP-A Produced by the AMBER Force Field Under GRO-MACS T-REMD

**Figure 2.**
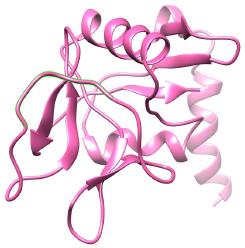
Model of SP-A Produced by the OPLS Force Field Under GRO-MACS T-REMD

**Figure 3.**
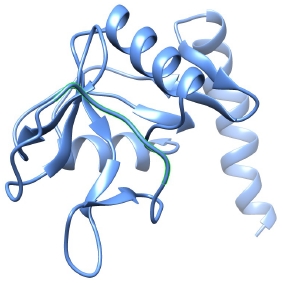
Model of SP-A Produced by the CHARMM Force Field Under GROMACS T-REMD

**Figure 4.**
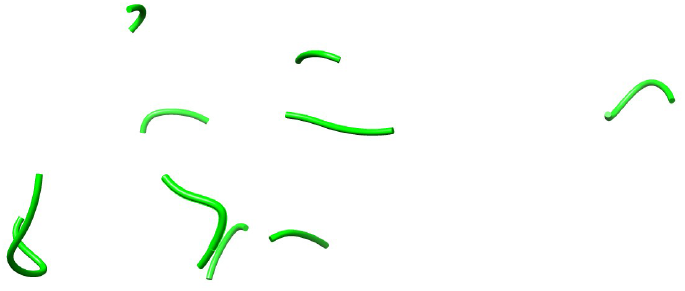
Model of SP-A Produced by Polar+ After Weak Binding Sites Were Cut

**Figure 5.**
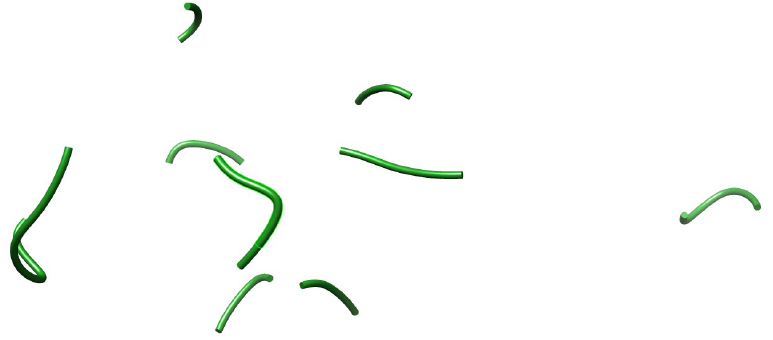
Model of SP-A Produced by the Goemans-Williamson Implementation After Weak Binding Sites Were Cut

#### 2. ZDOCK affinity values cannot be directly compared between graph-cut and T-REMD models

The complex with the highest ZDOCK conformation was the SP-A output of GROMACS using the AMBER force field and a model of SARS-CoV-2 spike protein without any modifications with the value of 2017.295. The lowest ZDOCK score observed was one of 448.202 for the complex consisting of Polar+ processed SP-A and Spike. However, due to the shape complementarity term of the ZDOCK algorithm (Ssc) creating a scalar based upon the degree of path matching between receptor and ligand structures and a lack of structural elements in models produced through graph cutting, the Ssc component for all Polar+/Goemans-Williamson models will be much lower than those produced by GROMACS, and cannot be directly compared.

#### 3. RMSD across ZDOCK conformations lower for graph cut, but likely inapplicable

The average RMSD across the top 2000 ZDOCK produced conformations of each model under consideration was calculated using ZDOCK output coordinate file. As a result, we found average RMSD values ranging from 219.7738 for the complex consisting of Polar+ processed SP-A and all atom, non processed Spike protein to 43.5559, for the complex of SP-A and Spike both processed via Polar+. Because the value of 219.7738 is relatively high considering the average atomic distance within SP-A and the SARS-CoV-2 spike, we hypothesize that this is not statistically significant configuration, as there is little convergence across the top models. However, having the top 2000 models produce an RMSD of 43.5559 is substantially lower than the next smallest value across the examined complexes, which is 133.9383 for the complex consisting from GROMACS OPLS force filed output SP-A and 6VSB Spike, showing that the model of SP-A and Spike both processed via Polar+ could be the most statistically significant configuration across the complexes under consideration. But, this is likely due to the fact that a model with fewer atoms has fewer possible configurational matches during binding, which would make the overall RMSD lower across graph-cut models.

### Concluding Remarks

Through this study, it was found that the operation of a quantum abstract machine (QAM) application of the Quantum Approximate Optimization Algorithm (QAOA) aimed at determining graph cuts within a protein structure in order to find top binding site candidates produced the same top binding sites and was faster than a QPU based version of itself, as well as faster than the Goemans-Williamson algorithm operated on the Bridges supercomputer, and the GROMACS T-REMD system on the JUWELS and Bridges supercomputer due to its superior computational complexity. Moreover, when used to provide binding results using the ZDOCK software, it produced results of chemical merit. However, the models produced by GROMACS for which the same binding test was completed had noticeably different binding results and configurations after this binding site determination step, binding directly to the SARS-CoV-2 Spike S1. For this reason, it is difficult to determine which model is superior without the completion of a biochemical assay to determine the actual method of interaction between the SARS-CoV-2 Spike and SP-A.

But, the computational improvements that quantum algorithms can provide in graph cutting is definitely promising in the quest to improve the computational time of complex analysis prior to protein-protein binding. Therefore, QAMs running quantum algorithms do seem to be an interesting area to investigate towards improvements of molecular dynamics simulations, especially with the addition of quantum specific state analysis through the determination of base and excitation energies using the variational quantum eigensolver (VQE) or similar [37] to provide improvements to quantum mechanics based models that can be completed in GROMACS today.

Lastly, this study did little to parameterize the qubits for the different thermal environments, which could have provided at par or higher accuracy in the type of modeling generated through graph cutting. Through the introduction of thermal relaxation models [32] to each bond site, the identification of qubits that best represent the electrostatic environment of the atom, the QPU can provide models that can represent thousands of time steps in real time due to the coherence timing of the qubits representing real, physical interactions, thus outperforming GROMACS on a supercomputer in terms of number of ns/day modeled.

## Acknowledgments

This study would like to give its greatest gratitude to Ken Hackworth and the team at Pittsburgh Supercomputing Center (PSC) and at XSEDE for awarding providing 1500 hours of compute time in order to complete the Goemans-Williamson portion of this study, Rigetti and Co for providing credits to use their quantum computing platform, Mark Skillbeck and Tom Lubowe of Rigetti for providing software troubleshooting help, the team at Forschungszentrum Jülich for providing computing time to complete the GROMACS portion of this study, Dr. Angela Haczku and her lab, including Angela Linderholm, Melissa Teuber, and Somy Cho, of the Pulmonary, Sleep, and Critical Care Divsion of the UC Davis Medical Center Department of Internal Medicine for providing insight into how to characterize surfactant proteins and its potential prevalence in SARS-CoV-2, and Kirk McGregor for providing insight into how to compare computational complexities among each of the software packages used and for editing support.

## Notes

### Competing Interest Statement

Samarth Sandeep is the CEO of Iff Technologies. Sona Aramyan is the CSO of Iff Technologies. Vaibhav Gupta is the CTO of Iff Technologies.

